# BREAKING THE TIGHT GENETIC LINKAGE BETWEEN THE *a1* AND *sh2* GENES LED TO THE DEVELOPMENT OF ANTHOCYANIN-RICH PURPLE-PERICARP SUPER-SWEETCORN

**DOI:** 10.1101/2022.07.13.499986

**Authors:** Apurba Anirban, Alice Hayward, Hung T Hong, Ardashir Kharabian Masouleh, Robert J Henry, Tim J O’Hare

## Abstract

The existence of purple-pericarp super-sweetcorn based on the most common supersweet mutation, *shrunken2* (*sh2*), has not been previously reported, partly due to its extremely tight genetic linkage to a non-functional anthocyanin biosynthesis gene, *anthocyaninless1* (*a1*). Generally, both aleurone- and pericarp-pigmented purple corn is starchy, the latter of which contains significantly higher anthocyanin compared to the former. The development of purple-pericarp super-sweetcorn is dependent on breaking the *a1-sh2* tight genetic linkage, which occurs at a very low frequency of <1 in 1000 meiotic crossovers. Here, to develop purple-pericarp super-sweetcorn, an initial cross between a male purple-pericarp maize (purple-round seed), ‘Costa Rica’ (*A1Sh2*.*A1Sh2*) and a female white *shrunken2* super-sweetcorn (white-shrunken seed), ‘Tims-white’ (*a1sh2*.*a1sh2*), was conducted. Subsequent self-pollination based on purple-pericarp-shrunken kernels identified a small frequency (0.08%) of initial heterozygous F3 segregants (*A1a1*.*sh2sh2*) producing a fully *sh2* cob with a purple-pericarp phenotype, enabled by breaking the close genetic linkage between the *a1* and *sh2* genes. Resulting rounds of self-pollination generated a F6 homozygous purple-pericarp super-sweetcorn (*A1A1*.*sh2sh2*) line, ‘Tim1’. Genome sequencing revealed a recombination break between the *a1* and *yz1* genes of the *a1-yz1-x1-sh2* multigenic interval, with the sequence pattern of ‘Tim1’ similar to the ‘Costa Rica’ parent, and after the linkage break, similar to the ‘Tims-white’ parent. The novel purple-pericarp super-sweetcorn produced a similar concentration of anthocyanin and sugar as in its purple-pericarp maize and white super-sweetcorn parents, respectively, potentially adding a broader range of health benefits than currently exists with standard yellow/white sweetcorn.

## Introduction

Unlike yellow or white super-sweetcorn, purple-pericarp super-sweetcorn based on the *shrunken2* gene mutation has not been reported previously. A principal reason for the absence of purple-pericarp super-sweetcorn is that an inactive allele of the anthocyanin biosynthesis pathway gene, *anthocyaninless1 (a1)*, is situated extremely close to the shrunken2 (*sh2)* supersweet mutation (Civardi et al., 1994), which provides the basis for the majority of super-sweetcorn globally (Revilla et al., 2021). The proximity of these genes, which are on chromosome 3, has been estimated as only <0.1 centi-Morgan (cM) (Civardi et al., 1994) with a physical distance of 140 kb (Yao et al., 2002). Therefore, simple crossing does not generate a normal meiotic segregation ratio of 3:1, but rather a low frequency crossover occurring at less than 1 in 1,000 plants. As a result, breaking the genetic linkage between these two genes is potentially challenging.

The pericarp of corn comprises the outer layers of the kernel/seed with four layers of cells, which is maternal tissue; whereas, the aleurone, which is situated just beneath the pericarp layer, is comprised of a single cell layer (Becraft and Yi, 2011). Pericarp-pigmented corn contains eight-times more anthocyanin than aleurone-pigmented corn (Paulsmeyer et al., 2017).

In purple corn (starchy maize), the dominant *A1* (*Anthocyanin1*) gene is closely linked (<0.1 cM) to the dominant allele of the *Sh2* gene (Kramer et al., 2015), and consequently the kernel of the corn is purple and round (starchy and non-sweet), and can easily be phenotyped by visual inspection. In *sh2* sweetcorn with the recessive *a1*, the kernels are shrunken when mature, and normally yellow or white depending on the underlying carotenoid pigmentation of the endosperm (O’Hare et al., 2015), in conjunction with colourless pericarp and aleurone layers. The absence of starch causes the kernel to adopt a shrunken appearance when it becomes fully mature and loses moisture. Anthocyanin is not formed in *sh2* sweetcorn, primarily due to the absence of a functional *A1* gene (Chhabra et al., 2019). Anthocyanin pigmentation in purple maize requires at least one copy of each of the dominant anthocyanin-biosynthesis genes (e.g., *A1*) to produce anthocyanin (Chaves-Silva et al., 2018).

An anthocyanin biosynthesis structural gene, *Pr1 (purple/red aleurone1)*, determines whether the anthocyanin-type will be purple (cyanidin) or red (pelargonidin) (Sharma et al., 2011). This gene is expressed in both the aleurone layer and pericarp, with its dominant form being responsible for cyanidin development and its recessive form for pelargonidin.

In addition, one regulatory gene from each of the Myb and bHLH transcription factor families is required (Hernandez et al., 2004; Petroni et al., 2014; Petroni and Tonelli, 2011) for anthocyanin biosynthesis. The Myb transcription factor *Pl1 (purple plant1)* is associated with light-independent pigment (purple) formation in the pericarp of the maize kernel, while *C1 (coloured aleurone1)* is responsible for aleurone-located anthocyanin development (Procissi et al., 1997). Similarly, the bHLH transcription factor *B1* (*plant colour1*) regulates plant colour, while *R1 (coloured1)* regulates aleurone as well as pericarp anthocyanin development (Chatham et al., 2019; Petroni et al., 2014).

Purple sweetcorn based on the *sugary1* (*su1*) gene mutation has been reported previously (Lago et al., 2014). However, sweetness enabled by the *sh2* mutation is more than two-fold that of the *su1* and eight-fold that of starchy corn (Feng et al., 2008). Although the aleurone-pigmented *sh2* sweetcorn has been reported previously (Civardi et al., 1994; Inplean et al., 2020; Yao et al., 2002), there has been no report of pericarp-pigmented sweetcorn based on the *sh2* mutation. Considering that the anthocyanin concentration of pericarp-pigmented corn is significantly higher than that of aleurone-pigmented corn (Luna-Vital et al., 2017), and the maternal nature of pericarp colour is not immediately transferable by pollen drift into adjacent yellow sweetcorn stands, the development of a purple-pericarp super-sweetcorn has both potential visual and agronomic benefits over aleurone-pigmented sweetcorn.

This study successfully developed purple-pericarp super-sweetcorn by breaking the close genetic linkage of *a1-sh2* to form *A1-sh2*. The recombination break point between the *a1* and *sh2* genes was demonstrated using genomic analysis. Finally, the anthocyanin and sugar concentrations of the developed purple-pericarp super-sweetcorn were found to be comparable to the purple maize and white *sh2* super*-*sweetcorn parents.

## Materials and methods

### Parental material and seedling establishment

For the development of purple-pericarp super-sweetcorn, a hand-pollinated initial cross was made between the white *shrunken2* super-sweetcorn ‘Tims-white’ (Fig 1a), as the female recipient, and the purple-pericarp Peruvian maize ‘Costa Rica’ (Fig 1b), as the pollen source. A reciprocal cross was also conducted to confirm pericarp tissue pigmentation was a maternal characteristic and not transferred by pollen. All plant materials were grown at the Gatton Research Facility, Department of Agriculture and Fisheries, Queensland, Australia from 2018 to 2021. Trials were conducted in isolation from other maize. Seed were directly sown, with plants separated by a 15 cm spacing and rows separated by 30 cm spacing. The soil was irrigated using 50 mm solid set pipes during initial plant establishment. Irrigation and fertilizers were applied during the plant’s life cycle through a trickle irrigation system. Similarly, pesticide was applied through the trickle irrigation system to protect the plants from soil-dwelling pests.

**Fig 1.**
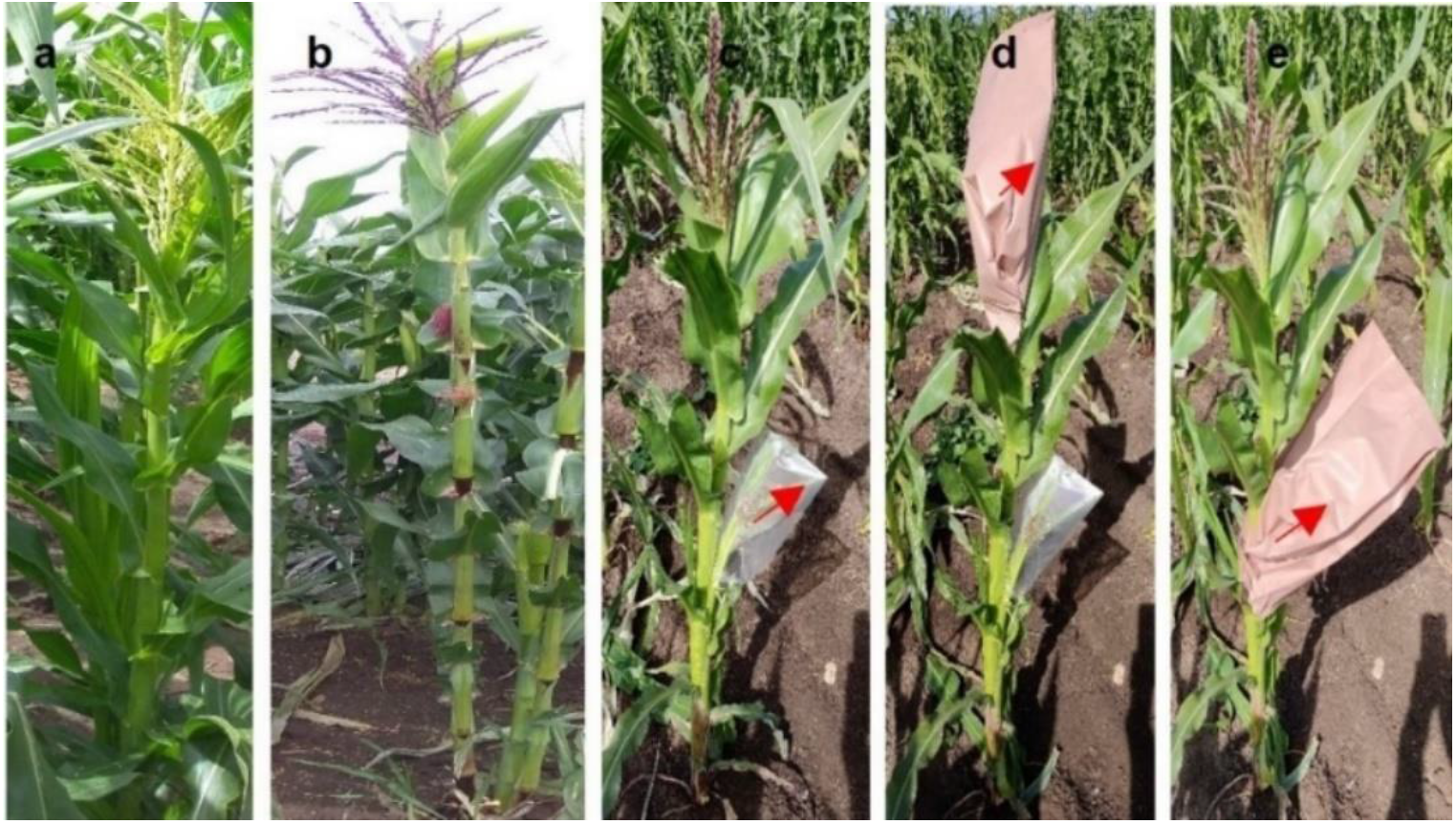
Parental plants: **sfiga)** white *shrunken2* super-sweetcorn accession, ‘Tims-white’, **b)** purple-pericarp Peruvian maize accession, ‘Costa Rica’. **Pollination and crossing: c)** covered ear, **d)** covered tassels (and ear), **e)** bagging after pollination.

### Pollination and crossing

Pollination was conducted under controlled conditions. Hand pollination was performed by initially covering the uppermost female ear with a clear plastic bag prior to silk emergence. After initial emergence of the silks, the silks were severed partway down the ear (approximately 3 cm back from the tip) to generate a uniform ‘brush’ of silks the next day to enable synchronised pollination. At the time of cutting the ear, the tassel from the male pollen donor was covered with a brown paper bag and secured with a paperclip. The following morning, the pollen bag was removed from the tassel, and the pollen from the bag applied to the emerged female silk ‘brush’. The pollinated ear was then covered with the paper bag and stapled in place to prevent further open pollination (Figs 1c, 1d & 1e).

### Self-pollination, plant emergence and phenological selection (phenotyping)

Fully mature F1 seeds generated from the crossed cob were harvested at maturity (50-60 days after pollination). Seeds were subsequently sown in separately marked rows, totalling 9 rows, each with 30 seeds per row. At flowering, silks were self-pollinated using pollen from the tassel of the same plant. The plants were then phenotyped for those that produced either white (no pericarp pigmentation) or purple-pericarp pigmented cobs (F2 generation). Those plants that produced white cobs were identified, but not used for further development.

From the purple-pericarp F2 cobs, kernels that exhibited a shrunken appearance when fully mature (approximately 25% of kernels per cob), were selected and pooled for each of the plant-rows from which they were obtained. Other phenological characteristics of the plants that produced purple-pericarp cobs were also documented, particularly anthocyanin production in other plant tissues. The F2 shrunken seeds were subsequently re-sown into a total of 92 rows, with 30 seeds sown per row to generate F3 plants.

At anthesis, plants were again self-pollinated, as described above. During growth, plants were visually phenotyped for evidence of anthocyanin production on the leaf-sheath, stem, leaf-auricles, silks and anthers. At full kernel maturity, all self-pollinated cobs were harvested, and the cob phenotype confirmed for kernel shape (shrunken or round) and pericarp colour (purple or white). The total number of plants, and those with purple-pericarp were counted in order to estimate the rate at which linkage breakage between *a1* and *sh2* occurred relative to the number of total plants.

All seeds exhibiting a purple-shrunken phenotype were harvested from those cobs exhibiting 100% purple-pericarp shrunken kernels. These were subsequently planted in individual rows (at least one row per purple-pericarp cob) and plants again self-pollinated at anthesis. The resulting cobs were phenotyped for pericarp colour, with a predicted segregation ratio of 3:1 purple-pericarp/white-pericarp cobs. The F4 purple-pericarp cobs were collected, and white-pericarp cobs discarded. Kernels from each F4 purple-pericarp cob were subsequently resown in individual rows. These were grown to maturity, self-pollinated, and the frequency of plants producing purple-pericarp cobs within a row recorded, with any row producing 100% purple-pericarp shrunken-kernel cobs considered homozygous for purple-pericarp super-sweetcorn (F5 & F6 cobs).

### Genomic analysis

DNA of 14-day-old leaf samples of Costa Rica’, ‘Tims-white’ and the developed purple-pericarp super-sweetcorn F6 line (‘Tim1’) was extracted according to a previous method by Vivekananda et al., (2018), with some modifications. The quality of the extracted DNA was assessed by spectrometry using the absorbance ratios of A260/280 nm and A260/230 nm (NanoDrop, Thermofisher Scientific). An absence of shearing of DNA and an A260/280 absorbance ratio of between 1.8-2.0, and a A260/230 absorbance ratio of between 2.0-2.2, were considered as high molecular weight DNA suitable for analysis. DNA re-sequencing (Illumina) was performed by Genewiz, China. Variants such as SNP (single nucleotide polymorphisms) and InDels (insertion, deletion mutations) in the *a1-sh2* interval were identified and compared with the publicly available annotated maize B73 (yellow starchy maize) (NAM5.0) reference genome (Woodhouse et al., 2021) using Qiagen CLC Genomic Workbench 21.0.4.

### Anthocyanin quantification

Cobs were harvested at the milk stage, 23 days after pollination (DAP). The kernels were removed manually from the cobs and stored at -20 °C. A row of kernels (about 15 kernels) was placed into milling vessels and dipped in liquid nitrogen for 3 min. The vessels were placed in a MM400 Retsch Mixer Mill (Haan, Germany) operated at 30 Hz for 60 s. A subsample of frozen powder was used for the determination of anthocyanin and sugar content. Anthocyanins were extracted from the frozen powder sample (∼1 g) following the method of Hong et al., (2020b). The anthocyanin analysis was determined using an ultra-high performance liquid chromatography–diode array detection-mass spectrometry (UHPLC-DAD-MS, Shimadzu, Kyoto, Japan) following the method of Hong et al., (2021).

### Sugar quantification

Sugar extraction of 23 DAP kernels was carried out as reported previously by Hong et al., (2021). Briefly, frozen powdered samples (0.5 g) were homogenized with 10 mL of aqueous methanol (70%). The mixtures were sonicated for 30 min at 50 °C and then shaken on a horizontal RP 1812 reciprocating shaker (Victor Harbor, SA, Australia) at 250 rpm for 10 min. Samples were then centrifuged at 4,000 rpm for 10 min at 50 °C. The supernatants were collected, and the pellets were re-extracted twice, following the same procedure. The sugar extracts were combined and diluted 20 times with aqueous acetonitrile (50%). The diluted sugar extracts were filtered through a 0.2 *µ*m hydrophilic PTFE syringe filter into HPLC vials for sugar analysis.

### Moisture content

Moisture content of kernels was conducted to ensure that harvested kernels were at a moisture content for optimal eating quality of *sh2* sweetcorn. Optimal moisture content should be within the range of 70-75% (Wann et al., 1971). Moisture content of harvested kernels was determined by taking three samples of each sweetcorn line, using the AOAC method 934.01 (AOAC, 1990) and the following formula:

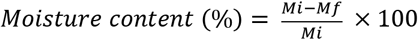

Here, Mi means initial weight and Mf means final weight of the powdered sweetcorn kernels. To get the final weight, the kernels were dried in a vacuum oven for 48 hours to remove moisture.

### Statistical analysis

One-way factorial analysis of variance (ANOVA) using the statistical software Prism (GraphPad Prism 9.3.1) was applied to assess variances and pairwise multiple comparisons. Fisher’s least significant difference (LSD, p < 0.05) was used to compare differences between means. Three replicates were taken from a cob, with 15 kernels comprising each replicate.

## Results

### Cross between purple maize and super-sweet parents to derive F1 seeds

The initial cross between white sweetcorn accession ‘Tims-white’ (Fig. 2a) with the purple-pericarp maize accession ‘Costa Rica’ (Fig 2b) yielded F1 cobs (*A1Sh2*.*a1sh2*) with white pericarp (Fig 2c), when the white sweetcorn was used as the female recipient. Approximately 50% of the kernels exhibited blue-aleurone pigmentation (Fig 2c). A concurrent reciprocal cross using ‘Costa Rica’ as the female and ‘Tims-white’ as the male generated round kernels with purple-pericarp (Fig 2d).

**Fig 2.**
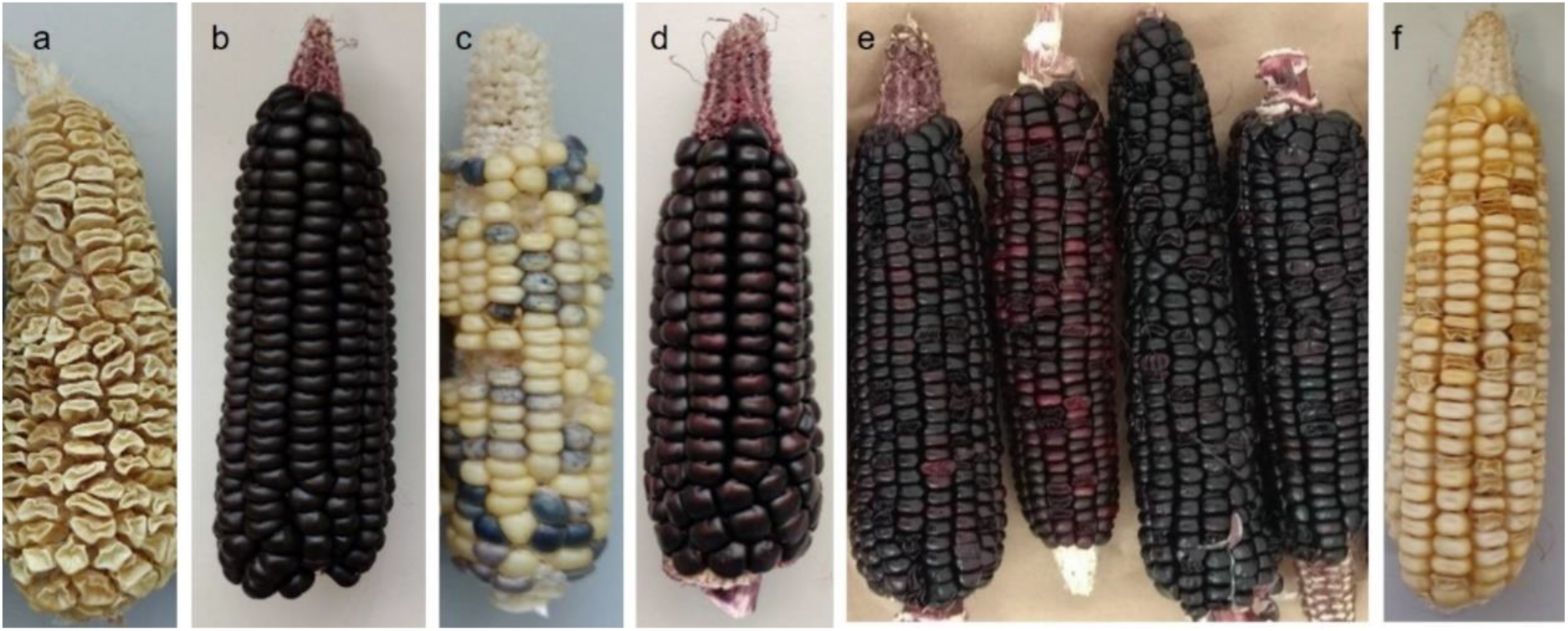
Parents and F1 progenies: **a)** white sweetcorn parent (‘Tims-white’-derived accession); **b)** Purple maize parent (‘Costa Rica’); **c)** F1 round kernels with white-pericarp, with or without blue-aleurone; **d)** reciprocal cross showing F1 round kernels with purple-pericarp. **F2 progenies**: **e)** cobs with purple-pericarp and reddish-purple-pericarp kernels with a 3:1 ratio of round to shrunken kernels; **f)** white cob with a 3:1 ratio of round to shrunken kernels.

### Production of F2 plants with purple-pericarp-shrunken seeds

F2 plants, obtained from the F1 seeds above, were grown to maturity, during which they were visually assessed for anthocyanin pigment synthesis in their cobs and various plant tissues. Out of 151 plants, a total of 119 plants were successfully self-pollinated. From an overall 119 self-pollinated cobs, the total number of cobs with purple kernels generated was 55 (45%), with the remaining cobs being white in colour. From the F2 cobs with purple kernels, some produced cobs with all purple-pericarp kernels, but in some, the kernel colour segregated into purple and reddish-purple or red pigmentation (Fig 2e).

Within all purple and white F2 cobs, as expected, the mature kernels were observed to segregate into 25% shrunken phenotype and 75% non-shrunken (round) (Figs 2e & 2f). From the purple-pericarp F2 cobs, from a total of 11,040 seeds, 2760 shrunken kernels were removed and subdivided into purple, reddish-purple and red shrunken kernels, respectively, for subsequent sowing. The white cobs were discarded.

### Linkage break at F3 generation and identification of purple-pericarp super-sweetcorn cobs

From the 2,760 self-pollinated F2 seeds sown, 1,532 germinated and produced cobs. Those plants that produced purple pigmentation in either the sheath-leaf, stem, auricle, anthers and silks (Fig 3a-e) were selected for controlled self-pollination. Consequently, 15 plants were observed to produce purple colouration in at least one of the above tissues during the growth period.

**Fig 3.**
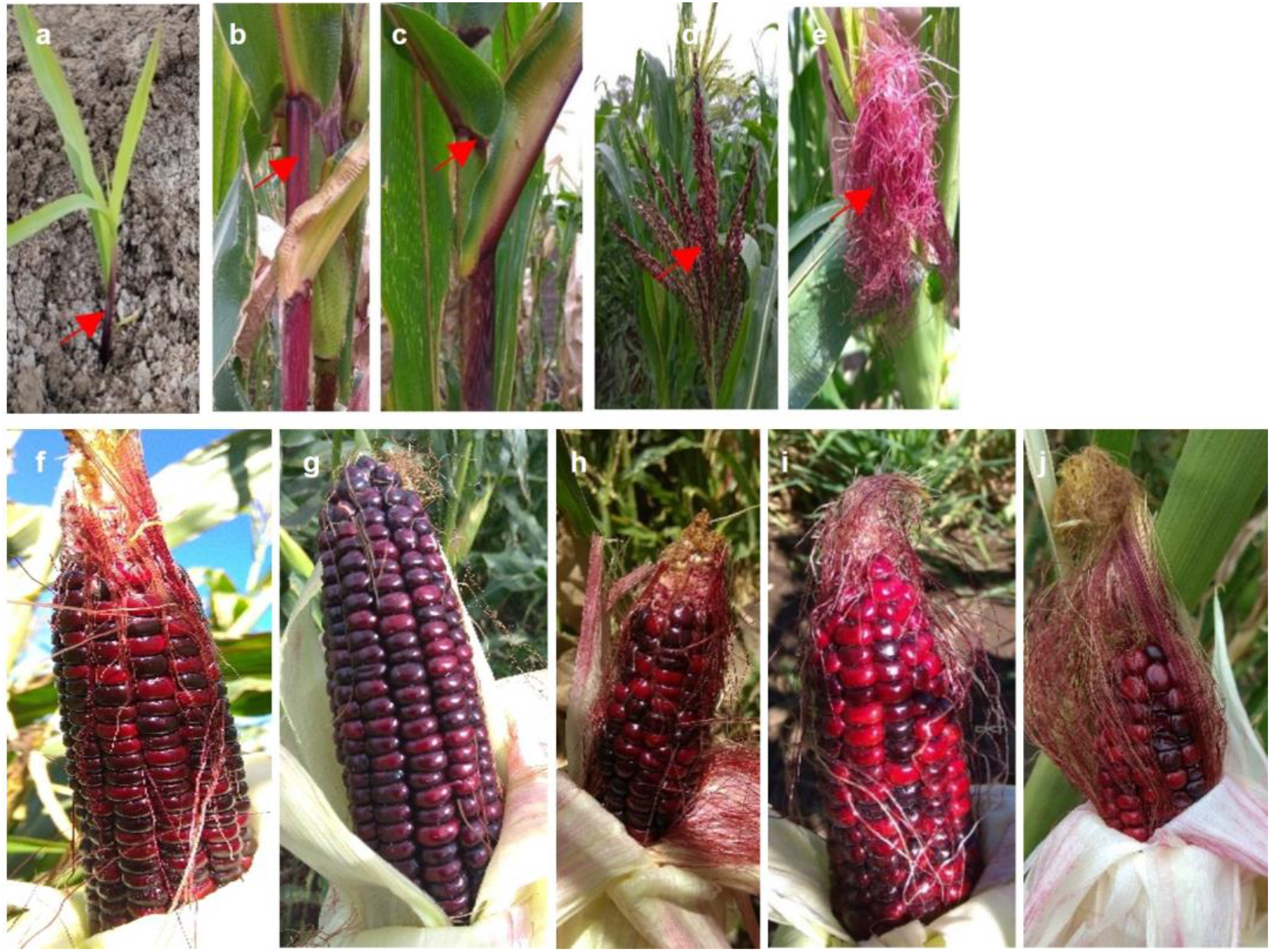
Purple pigmentation in different tissues of F3 plants at different stages of plant development: a) sheath-leaf, b) stem, c) auricle, d) anthers, and e) silks. **Developed purple F3 cobs:** ‘Tim1’ to ‘Tim5’ (f-j) at milk/eating stage, prior to exhibiting the mature shrunken phenotype.

During kernel development, three of the above 15 plants with purple pigmentation produced cobs with mainly purple-pericarp kernels (Figs 3f, 3g & 3h). In addition, two cobs were also identified to have some purple-pericarp and maximum red-pericarp kernels (Figs 3i & 3j). Each of these five cobs were confirmed to develop a shrunken kernel phenotype at full kernel maturity (60 DAP). Other plants within the 15 were observed to produce purple-pericarp, but at maturity the kernels exhibited a non-shrunken (round) phenotype and were not considered for further study.

The five identified purple-pericarp shrunken-kernel super-sweetcorn accessions were named as ‘Tim1’, ‘Tim2’, ‘Tim3’, ‘Tim4’, and ‘Tim5’.

### Germination analysis and frequency of linkage breaking

As referred to above, a total of 1,532 seeds were germinated from 2,760 F2 purple-pericarp shrunken seeds planted. As a result, the percentage of germination of the F3 plants was 56%. Therefore, from a total 11,040 original F2 seeds (including both round and shrunken), it was extrapolated that total germination would have been 6,128.

From the 1,532 F3 plants, five plants were observed to produce cobs with purple-pericarp shrunken kernels (*A1a1*.*sh2sh2*), equating to a linkage break having occurred. Among these, four (‘Tim1’, ‘Tim2’, ‘Tim4’ and ‘Tim5’) were germinated from F2 seeds that had been further classified as purple seeds, while the other (‘Tim3’) was from seed classified as being reddish-purple. No linkage break was observed from plants with originally red seeds.

Based on the production of five plants with a purple-pericarp and shrunken phenotype from the original 6,128 viable F2 seeds (based on extrapolated germination viability) a linkage break between *a1* and *sh2* was estimated to occur 1 in every 1,226 plants (5 in 6,128 plants), equating to 0.08 centi-Morgan (cM) genetic distance between the *a1* and *sh2* genes.

### F4 super-sweetcorn cob development and segregation of purple-pericarp and white-pericarp cobs

In the F3 generation, five purple-pericarp shrunken kernel segregants were identified, each putatively with the *A1a1*.*sh2sh2* genotype, heterozygous for *A1*.*a1* (Figs 3f-j) as indicated in Table 1.

**Table 1.**
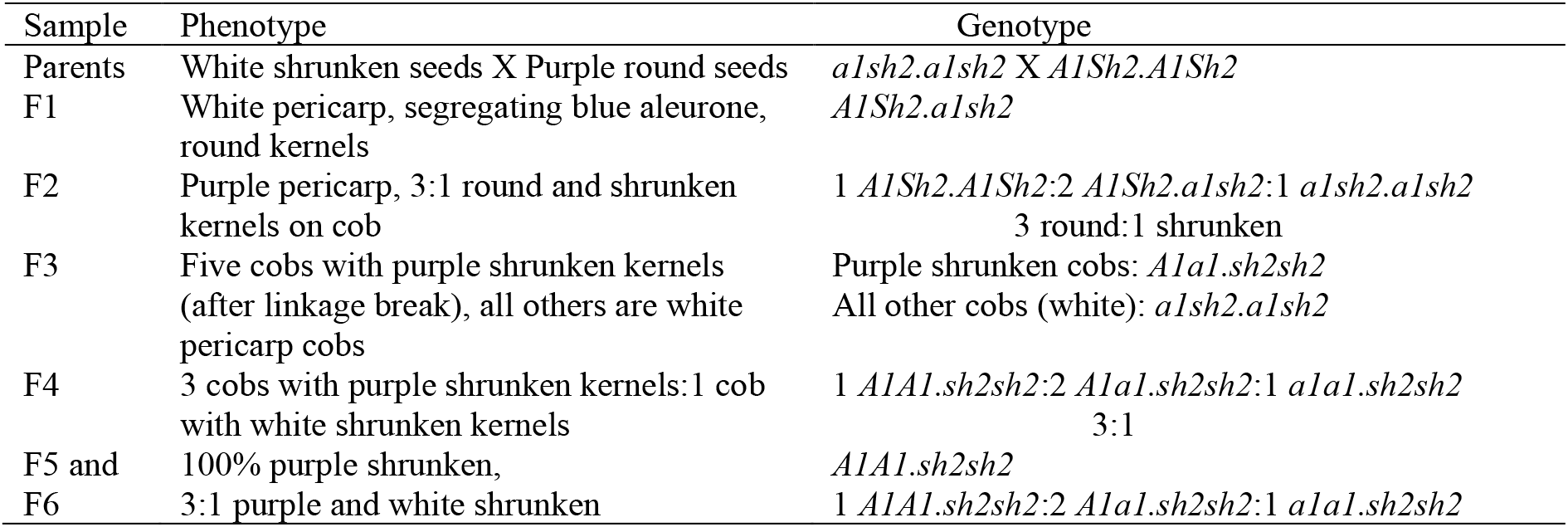
Expected phenotypes and genotypes of each experiment

In the subsequent F4 generation, kernels derived from F3 ‘Tim1’ line segregated as 3:1 purple-pericarp and white-pericarp cobs, confirming the heterozygous genotype (*A1a1*.*sh2sh2*) of the ‘Tim1’ parent (F3) plant. ‘Tim5’ also yielded a 3:1 cob-ratio with purple- and white-pericarp, respectively, however plants from the red seeds of ‘Tim5’ produced only white cobs and green plants with no sign of purple pigmentation in any tissue. ‘Tim2’ produced some plants with purple tissues as well as underlying cobs with purple-pericarp, however the germination was low due to fungal infection. Germination of ‘Tim3’ kernels was also very low and produced only green plants with no purple pigmentation, so was discarded for the next experiment. ‘Tim4’, however, presented a reverse scenario, in which 1:3 plants with purple-pericarp and white-pericarp cobs were obtained.

### Development of F5 super-sweetcorn cob homozygous for anthocyanin biosynthesis gene

In the F5 generation, all progeny generated from seeds of one cob of F4 plant derived from ‘Tim1’produced 100% purple-pericarp cobs, indicating a homozygous purple-pericarp sweetcorn genotype (*A1A1*.*sh2sh2*). In contrast to the above, one cob of another F4 plant derived from ‘Tim1’, as well as a cob of F4 plants derived from ‘Tim4’, and two cobs of F4 plants derived from ‘Tim5’, gave rise to a 3:1 ratio of purple-pericarp and white-pericarp cobs, respectively, confirming their heterozygosity (*A1a1*.*sh2sh2*). In contrast, red seeds of ‘Tim5’ produced all green plants, as in the previous experiment. Seeds from one cob of F4 plants derived from ‘Tim2’ and a cob of F4 plants derived from ‘Tim4’, produced a 2:1 ratio of purple-pericarp to white-pericarp cobs.

### Development of F6 cobs

In the subsequent F6 generation, seeds from the two F5 ‘Tim1’ progeny above produced 100% of plants with purple-pericarp cobs, confirming their homozygous nature (*A1A1*.*sh2sh2*). The remaining ‘Tim’ lines (‘Tim2’, ‘Tim4’ and ‘Tim5’), as well as plants generated from the third cob of ‘Tim1’, produced 3:1 purple-pericarp and white-pericarp cobs, confirming their segregating nature, regarding the structural anthocyanin gene, *A1*.

### Identification of the recombination break site between *a1* and *sh2* genes by genomic analysis

The B73 maize reference *a1* and *sh2* genes are 4,826 bp (221,778,277-221,783,103 bp) and 14,492 bp (221,890,353-221,904,845 bp) (Woodhouse et al., 2021) (Fig 4a) long on chromosome 3. There are two other genes (*yz1, x1*) situated within the *A1* and *sh2* interval (Yao et al., 2002) (Fig 4a). The linkage break formed from the meiotic recombination of purple-round ‘Costa Rica’ (*A1Sh2*.*A1Sh2*) and white-shrunken ‘Tims-white’ (*a1sh2*.*a1sh2*) parents is responsible for the presence of the dominant *A1* allele and the recessive *sh2* allele in the developed purple-pericarp super-sweetcorn line, ‘Tim1’ (Fig 4b).

**Fig 4.**
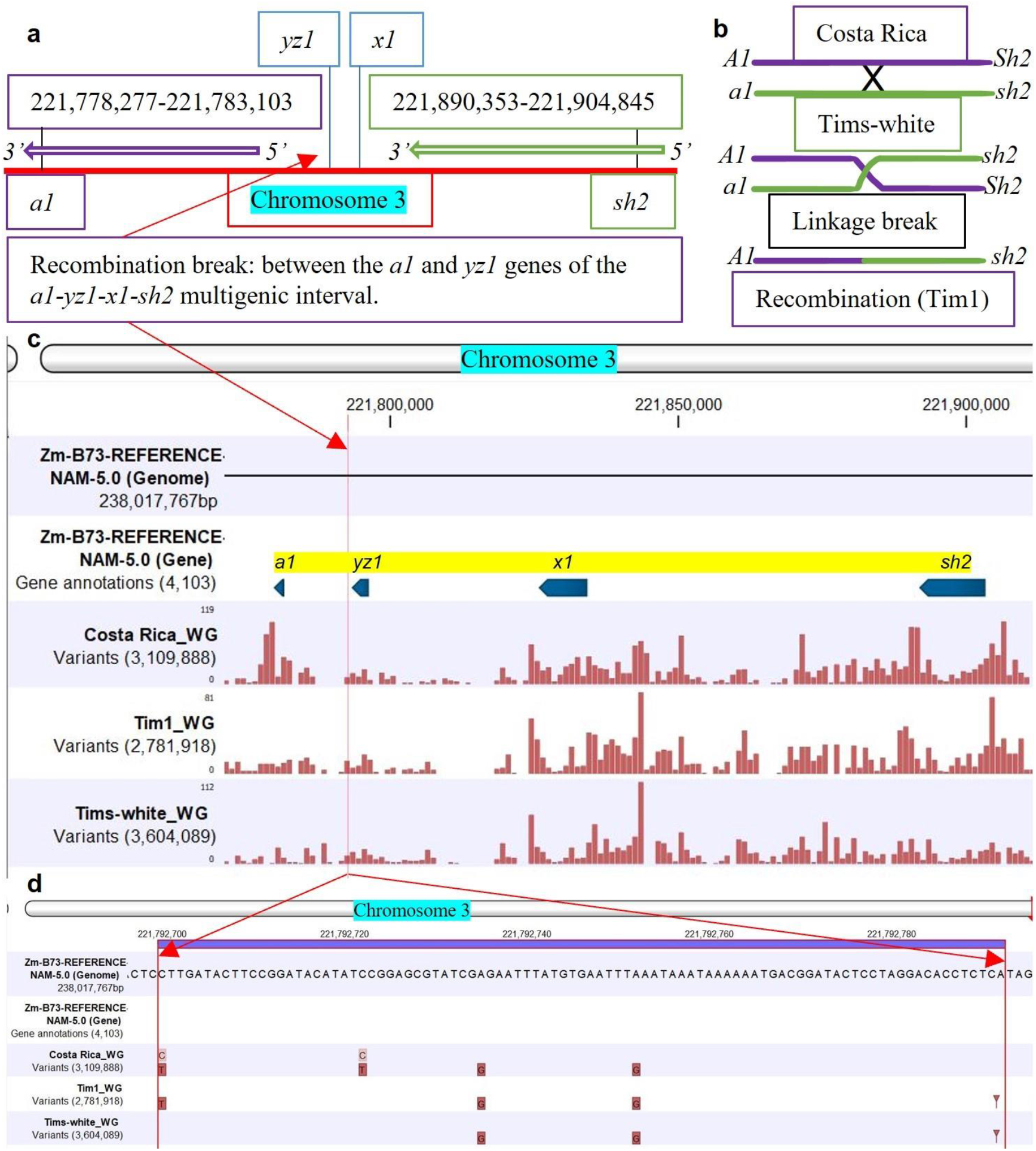
Linkage break and recombination: **a)** length of *a1* and *sh2* genes on chromosome 3, along with the position of *yz1* and *x1* genes; **b)** linkage and recombination between *A1* and *sh2* alleles; **c)** mapping of the whole genome (WG) of ‘Costa Rica, ‘Tim1’ and ‘Tims-white’ lines against the reference B73 genome, with four genes *(a1, yz1, x1* and *sh2*) within the *a1-sh2* interval annotated (blue arrow), red bar indicates variants, with height of bar meaning read coverage; **d)** red arrows showing putative linkage break and recombination site between *a1* and *sh2* genes of the developed F6 line, ‘Tim1’.

The recombination site between the *a1* and *sh2* genes was identified as being between the *a1* and *yz1* genes (Fig 4c) of the *a1-sh2* interval. Linkage break was observed between 221,792,700 bp and 221,792,792 bp (Fig 4d) of the reference genome. Therefore, it can be concluded that this site may be considered the putative linkage break site between the *a1* and *sh2* genes.

### Anthocyanin quantification of sweetcorn eating-stage kernels (23 DAP) of parental lines and the developed F5 line, ‘Tim1’

Cyanidin-, peonidin- and pelargonidin-based anthocyanin compounds of ‘Costa Rica’, ‘Tim1’ and ‘Tims-white’ were analysed by UHPLC-DAD. The developed purple sweetcorn F5 line, ‘Tim1’ (140.5 mg/100g FW, fresh weight) and ‘Costa Rica’ (142.2 mg/100g FW) parental line at 23 DAP possessed similar total anthocyanin concentrations (not significantly different, P<0.005) in their kernels (Fig 5a). In contrast, the white sweetcorn parent did not have any detectable anthocyanin.

**Fig 5.**
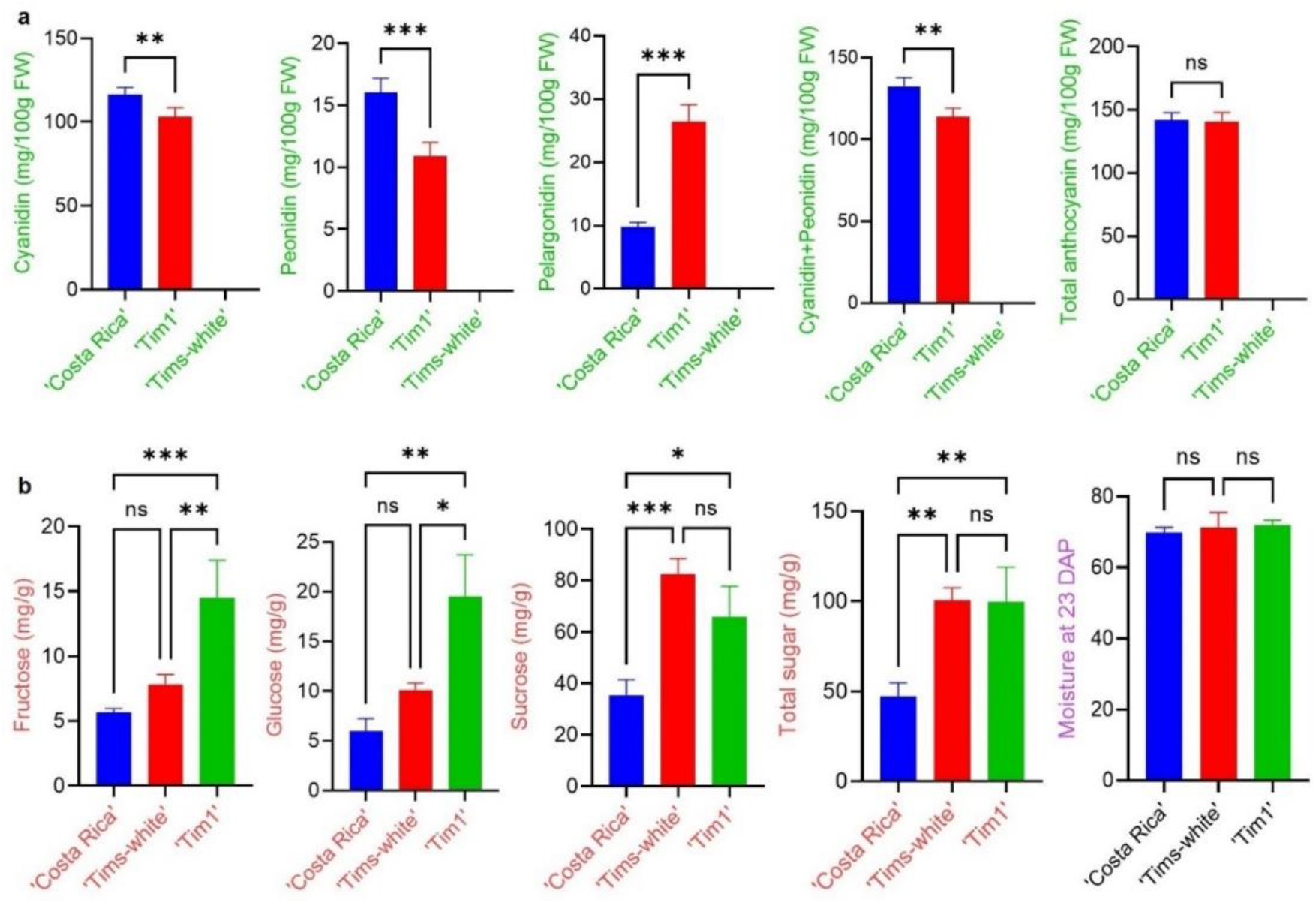
One-way factorial analysis of variance (ANOVA) using Fisher’s Least significant difference (LSD) test for anthocyanin and sugar concentrations. Multiple Comparison showing ns=non-significant, *= significant at P <0.005, **= significant at P <0.002, and ***= significant at P < 0.001. **a)** anthocyanin profiles of ‘Costa Rica’ and ‘Tim1’ kernels at 23 days after pollination. **b)** sugar profiles of ‘Costa Rica’, ‘Tims-white’ and ‘Tim1’ kernels at 23 DAP.

In both ‘Costa Rica’ and ‘Tim1’, cyanidin-based anthocyanins constituted the main proportion of anthocyanins. Although there was no significant difference in total anthocyanin concentration between ‘Tim1’ and ‘Costa Rica’, however, the ‘Costa Rica’ parent had a significantly higher proportion of cyanidin-based anthocyanins (82%) than ‘Tim1’ (73%), and the red pigment (pelargnidin-based glucoside) accounted for 7% and 19%, respectively. Peonidin-based anthocyanins also followed a similar trend to cyanidin-based anthocyanins, with 11- and 8% in ‘Costa Rica’ and ‘Tim1’, respectively, although peonidin generally accounted for only a small proportion of the total anthocyanin (Fig 5a).

### Sugar quantification of sweetcorn eating-stage kernels (23 DAP) of parental lines and the developed F5 line, ‘Tim1’

Sugars in ‘Costa Rica’, the F5 line ‘Tim1’ and ‘Tims-white’ were analysed by UHPLC-MS/MS. The constituent sugars detected consisted of fructose, glucose, and sucrose in all three lines, with sucrose being the principal sugar present (Fig 5b). Total sugar concentration was observed to be highest (100.5 mg/g FW, fresh weight) in the ‘Tims-white’ parental line and the F5 line ‘Tim1’ line (99.8 mg/g), with no significant difference between them at 23 DAP (Fig. 5b). By contrast, the purple maize, ‘Costa Rica’ produced a significantly lower (P<0.005) total sugar concentration (47.3 mg/g).

As with total sugar concentration, a similar pattern was observed with sucrose concentration, the principal sugar component (Fig 5b). Again, a significantly lower concentration of sucrose was observed for the ‘Costa Rica’ parent (35.57 mg/g) compared to the ‘Tims-white’ (82.57 mg/g) and ‘Tim1’ (65.85 mg/g) (Fig 5b). Although there was a non-significant trend (P<0.005) for sucrose concentration to be slightly lower in ‘Tim1’, the monosaccharide sugar constituents, glucose and fructose, were observed to be significantly higher than its ‘Tims-white’ parent (Fig 5b).

A similar moisture content (70-72%) was observed in all three lines at 23 DAP (Fig 5b), indicating that all three lines were at a sweetcorn eating-stage. Furthermore, observed differences in sugar concentration were not due to a difference in dry matter content between the three lines.

## Discussion

### Breaking the *a1-sh2* genetic linkage to develop heterozygous purple-pericarp super-sweetcorn

The present study was successful in breaking the close genetic linkage between the *a1* and s*h2* genes in a purple-pericarp maize, enabling the coexistence of a functional anthocyanin biosynthesis allele (*A1*) with the recessive supersweet allele, *shrunken2* (*sh2*). Although this has been previously achieved in aleurone-pigmented corn (Civardi et al., 1994, Yao et al., 2002), this is the first report of this occurrence in pericarp-pigmented maize. The potential benefit of a pericarp-pigmented sweetcorn over aleurone-pigmented sweetcorn is twofold. Firstly, the anthocyanin concentration is significantly greater in pericarp-pigmented corn, increasing visual intensity and increasing the concentration of a known phytonutrient (Hong et al., 2020a). Secondly, as the pericarp is maternal tissue, and only expressed by the maternal plant, it is not able to be passed across to adjacent stands of yellow sweetcorn, as is the case with ‘spotting’ caused by aleurone-pigmented sweetcorn (Carey et al., 2004).

In the current study, the initial cross between ‘Tims-white’, a white shrunken sweetcorn (*a1sh2*.*a1sh2*), and ‘Costa Rica’, a purple round (starchy) maize (*A1Sh2*.*A1Sh2*), produced F1 cob (*A1Sh2*.*a1sh2*) with white starchy (round) kernels and some kernels with some aleurone-based anthocyanin pigmentation (Fig 2c).

It is inferred that ‘Costa Rica’ parent may have been heterozygous (*Cc*) for the *C1* (*coloured aleurone1*) transcription factor gene, which is required for aleurone-based anthocyanin pigmentation (Procissi et al., 1997). This would potentially explain the segregating nature of aleurone colour in the cob (Fig 2c). As ‘Costa Rica’ of current study is based on a purple maize landrace, it has not been deliberately inbred, and therefore segregation of some genes, particularly those masked by overlying pericarp-pigmentation, is a likely probability. It is important however, if pollen contamination (spotting) is to be avoided in adjacent stands of yellow sweetcorn, that selection is made of lines that possess pericarp pigmentation, but lack aleurone pigmentation.

Reciprocal cross between ‘Costa Rica’ and ‘Tims-white’ produced purple-pericarp cob with round kernels (Fig 2d). The result of the reciprocal cross, however confirmed that the pericarp pigmentation is the same as the mother, and is therefore from a maternal trait/tissue as previously reported by Ron-Parra et al., (2016).

Interestingly, although pericarp tissue is maternal tissue, and therefore all kernels on the same cobs should be coloured the same (e.g., purple), it was apparent that there was variation in kernel colour between purple, reddish-purple and red within some of the F2 pericarp-pigmented cobs (Fig 2e). As pericarp tissue is maternal, the pericarp of all kernels on a single cob shares the same genotype as the mother plant. As such, the colour differences most likely came from underlying variation in aleurone colour, as absence of colour in the aleurone will make the overlying pericarp above look less purple, especially at sweetcorn eating stage, when anthocyanin is still accumulating (Hong et al., 2021). The reason behind this may be partly due to the absence or presence of aleurone pigmentation, but could also be due to inter-kernel differences in aleurone anthocyanin profile.

In maize and sweetcorn, the *Pr1 (purple/red aleurone1)* gene in the anthocyanin biosynthesis pathway regulates the direction of biosynthesis towards cyanidin or pelargonidin (Sharma et al., 2011). It may be inferred that the *Pr1* gene in the original purple ‘Costa Rica’ parent was homozygous dominant (*Pr1Pr1*), and the ‘Tims-white’ parent was homozygous recessive (*pr1pr1*), and after crossing (F1) they produced the heterozygous (*Pr1pr1*) form of *Pr1*. Once self-pollinated to form kernels in the F2 generation, *Pr1pr1* would have segregated into three allele combinations. It is probable that the aleurone of the darker purple F2 kernels have dominant homozygous (*Pr1Pr1)*, the reddish-purple seeds have heterozygous (*Pr1pr1*), and red seeds are homozygous recessive (*pr1pr1*). Consequently, whereas the pericarp be consistent in its anthocyanin profile across the cob, the underlying aleurone will vary in its profile, with the combination of pericarp and aleurone pigmentation combining to provide the observed kernel colour.

Kernels of the F2 purple-pericarp cobs segregated in a 3:1 ratio of round to shrunken kernels, as expected (Fig 2e). As the shrunken phenotype is only formed in relation to the homozygous recessive genotype *sh2sh2*, only the purple shrunken kernels were separated for the subsequent field experiment, as the shrunken phenotype was now fixed (*i*.*e*., homozygous). Generally, the recessive *sh2* allele is associated with a non-functional *a1* allele of the *anthocyanininless1* gene (Kramer et al., 2015), but breakage of this linkage between *a1* and *sh2* has been reported previously in aleurone-pigmented maize (Yao et al., 2002), such that an *A1a1*.*sh2sh2* genotype should occur at a frequency of <1 in 1000 crossover events (Civardi et al., 1994). Consistent with this, in this current study, a total of 2,760 shrunken kernels were collected and planted to enable this segregation. It is also worth mentioning that unlike pericarp-pigmented maize, different transcription factor genes are responsible for the activation of anthocyanin biosynthesis structural gene in aleurone-pigmented maize (Chatham et al., 2019; Petroni et al., 2014).

In the third field experiment, out of 15 plants exhibiting anthocyanin pigmentation in vegetative tissues, five purple-pericarp super-sweetcorn lines were subsequently identified (Fig 3). Based on the crosses, each of the five generated lines would have possessed a *A1a1*.*sh2sh2* genotype, as indicated in Table 1. The ten other lines producing anthocyanin were observed to have round kernels also produced on the cob. This indicated that the genotype was most likely *A1Sh2*.*a1sh2*, due to pollen contamination from a non-sweet purple-pericarp plant. As this trial was conducted in isolation from other maize, the origin of contaminating pollen may have possibly happened through mis-phenotyping of shrunken kernels during seed selection.

The genetic distance between the *a1* and *sh2* genes of the purple-pericarp super-sweetcorn developed in the above field trial was calculated to be 0.08 cM. This was comparable to previous research with aleurone-pigmented maize, in which Civardi et al., (1994) calculated a 0.09 cM genetic distance, and Yao et al., (2002) calculated a 0.07 cM genetic distance between the *a1* and *sh2* genes.

### Development of homozygous (fixed) purple-pericarp super-sweetcorn line

As the linkage break was only predicted to occur in one of the two copies of the *a1-sh2* allele association (i.e., *A1a1*.*sh2sh2*), an F4 generation was produced to confirm this, and also for the subsequent development of a homozygous genotype (i.e., *A1A1*.*sh2sh2*). F4 progenies of ‘Tim1’ and ‘Tim5’ produced a 3:1 ratio of cobs with purple and white pericarp kernels, however red seeds of ‘Tim5’ produced only white cobs, may be due to the absence of a functional anthocyanin biosynthesis gene, such as *A1*. ‘Tim2’ produced only a few plants with purple tissues due to the low germination rate caused by fungal infection, as reported in sweetcorn (O’Hare et al., 2015). As a result, the determination of identifying whether it was homozygous for the anthocyanin biosynthesis gene (*A1A1*) was achieved in a subsequent experiment. Very low germination was also observed in the ‘Tim3’ line, which only produced green plants, indicating an absence of a functional *A1* gene, and therefore was not considered for subsequent re-planting. The green plants germinated may have come from the recessive genotype of 3:1 segregation of F3 progenies (*i*.*e*., *a1a1*.*sh2sh2*).

Theoretically, all of the developed genotypes of the F4 generation should produce 3:1 cobs with purple- and white-kernels, however ‘Tim4’ produced 1:3, possibly due to the segregating nature of the anthocyanin biosynthesis structural (*A1*) as well as potential segregation of the regulatory genes, *Pl1* and *R1*. However, it may also have been possible that the low number of seeds used may have hampered the calculation of an accurate segregation ratio. Consequently, further investigation by subsequent field experiments was necessary.

In the fifth and sixth field experiments, a homozygous purple-pericarp super-sweetcorn line (‘Tim1’) was developed, theoretically with two copies of the dominant anthocyanin biosynthesis structural allele, *A1*, in conjunction with two copies of the recessive supersweet allele, *sh2*. This was based on the absence of any plants lacking purple-pericarp cobs. As such, the current study is apparently the first report of breaking the close genetic linkage between the *a1* and *sh2* genes in a purple-pericarp super-sweetcorn.

### Genomic analysis identifying the recombination break site between *a1* and *sh2* genes

The purple-pericarp super-sweetcorn line ‘Tim1’ was developed by breaking the tight genetic linkage between the *a1* and *sh2* genes, which enabled the recombination of a functional *A1* allele from the ‘Costa Rica’ parent with the recessive *sh2* supersweet allele from the ‘Tims-white’ parent leading to the formation of an initial heterozygous (F3) and subsequent homozygous (F6) line, ‘Tim1’ (*A1A1*.*sh2sh2*) (Fig 4b).

There are some common variants in both ‘Costa Rica’ and ‘Tim1’ before the linkage break (in the *a1* gene region) and some common variants in both ‘Tims-white’ and ‘Tim1’ after the linkage break (in the *sh2* gene region) (Fig 4c). Recombination was identified as occurring between the *a1* and *yz1* genes (Figs 4c & 4d) within the *a1-sh2* interval. Therefore, this site might be considered as defining the putative linkage break site of the *a1* and *sh2* genes, that allowed recombination of the *A1* and *sh2* alleles in ‘Tim1’. An earlier study by Yao et al. (2002) with meiotic recombination across the *a1-sh2* interval also found several recombination break points between the *a1* and *yz1* genes when coloured (*A1-LC*) and colourless (*a1::rdt*) alleles of maize were used.

### Anthocyanin profiling

Purple kernel pigmentation can be due to either pigmentation of the pericarp tissue, which is maternal (Ron-Parra et al., 2016), or the aleurone, which is triploid and therefore influenced by the pollen source as well as the parent plant (Ma et al., 2004). The pericarp comprises the outer layers of the kernel, about four layers of maternal cell tissue; whereas the triploid aleurone is situated just beneath the pericarp layer, usually consisting of a single layer cell (Becraft and Yi, 2011; Paulsmeyer et al., 2017). As such, pericarp-pigmented corn contains a higher anthocyanin content than aleurone-pigmented corn (Luna-Vital et al., 2017).

Sweetcorn, as a horticultural product is harvested usually at about 20-28 DAP, when the kernels are still physiologically immature (Hong et al., 2020a; Khanduri et al., 2011; O’Hare et al., 2015). However, starchy maize, which is used as flour (Lago et al., 2012) is harvested usually physiological maturity stage. As such, information on anthocyanin at purple starchy maize at eating stage is lacking. However, in a previous study by Hong et al., (2020a) with *brittle1* purple sweetcorn, total anthocyanin at 26 DAP was 47.77 mg/100g fresh weight and at increased maturity (36 DAP) was 179 mg/100g dry weight The total anthocyanin concentration of the developed line ‘Tim1’ at eating stage (23 DAP) found in this current research (140.51 mg/100g FW) (Fig 5a) was significantly higher than anthocyanin found in other crops, for example, coloured strawberries (60 mg/100g FW) (Aaby et al., 2012), red plums (30.1 mg/100g FW) (Proteggente et al., 2002), and red currants (12.8 mg/100g FW) (Wu et al., 2006), in which anthocyanins were not as high as in the purple-pericarp super-sweetcorn line, ‘Tim1’.

In the current study, the absence of anthocyanin in the *shrunken2* ‘Tims-white’ parent indicated an absence of a functional anthocyanin biosynthesis gene (e.g., *A1*). In addition, although there is a significant change in the proportion of anthocyanin components, the purple pigment (cyanidin-based and peonidin-based glucoside) in ‘Costa Rica’ parent and ‘Tim1’ is significantly much higher than the red pigment (pelargonidin-based glucoside). Previous study by Petrussa et al., (2013) indicated that peonidin is a methylated form of cyanidin and both cyanidin and peonidin form a purple colour. Therefore, the purple is the pigment appearing in both ‘Costa Rica’ and ‘Tim1’ pericarp-kernel lines studied.

As anthocyanin has been associated with a range of health benefits such as cancer chemoprevention, inhibiting colorectal carcinogenesis, lowering the body weight and reducing the risk of colon cancer as well as oxidative stress in rat, mice and human, respectively (Hagiwara et al., 2001; Hou et al., 2004; Miranda-Rottmann et al., 2002; Wu et al., 2017), and that anthocyanin is not normally associated with yellow/ white sweetcorn, the high concentration of anthocyanin observed in the current study indicates that these purple-pericarp super-sweetcorn lines could potentially present different health benefits than the yellow/white sweetcorn.

### Sugar profiling

The shrunken phenotype of sweetcorn arises from an absence of starch as the kernel matures. ADP-glucose pyrophosphorylase is the main enzyme in starch biosynthesis, however in *sh2* endosperms, ADP-glucose pyrophosphorylase activity is absent (Doehlert and Kuo, 1990), and hence it stops conversion of sugar to starch. The absence of starch is the cause of the shrunken appearance of the phenotype at full kernel maturity. The *sh2* is a recessive gene mutation and two copies of this gene is required to develop a *shrunken2* phenotype (Ruanjaichon et al., 2021).

In this study, moisture content of ‘Tims-white’ and ‘Tim1’ lines at 23 DAP was between 71-72%. Similarly, in a previous study by Wong et al., (1994) with different *sh2* sweetcorn varieties, moisture content was reported within a similar range of 71-73% at 23 DAP. It indicated that 23 DAP was within the sweetcorn eating stage range based on expected moisture content.

The total sugar concentration at 23 DAP of the purple-sweetcorn progeny, ‘Tim1’ was, not significantly different to the white-sweetcorn parent, ‘Tims-white’. However, the proportion of individual sugars was different (Fig 5b), with the ‘Tim1’ line expressing significantly greater fructose and glucose concentrations than its parent, ‘Tims-white’. Interestingly, sucrose concentration was not significantly different, and, as this constituted the main sugar present, also explain why total sugar concentration was also not significantly different between these lines. The principal sugar was identified as sucrose in both the parents and the progeny, which is in agreement with earlier studies based on *brittle1* and *sugary1* sweetcorn varieties (Hong et al., 2021; Vries and Tracy, 2016).

Total sugar concentration observed in the developed ‘Tim1’ purple and ‘Tims-white’ super-sweetcorn lines were more than two-fold that of the purple maize parent, ‘Costa Rica’. It confirms that due to the presence of dominant *Sh2* allele, the conversion of sugar to starch occurred in ‘Costa Rica’. Therefore, it has less sugar at milk stage, and all of them should be converted to starch at mature stage. This result supports previous study with waxy corn, in which no sugar was found at full kernel maturity (40 DAP) (Simla et al., 2010). By contrast, due to the presence of the recessive *sh2* allele, conversion of sugar to starch is hampered and more sugar is produced in the ‘Tim1’ purple and ‘Tims-white’ super-sweetcorn lines.

## Acknowledgements

This article is the part of PhD research of Apurba Anirban. AA is grateful to the University of Queensland for providing a PhD Fellowship.

## Author contributions

AA performed field experiments, DNA extraction, genomic analysis, anthocyanin and sugar extractions, and numerical calculations. AA performed field experiments with the help of TJO, DNA extraction with the help of AH and genomic analysis with the help of AH, AKM and RH. HTH analysed the HPLC data related to anthocyanin and sugar, and AA performed the statistical analysis. AA composed the manuscript. TJO supervised the PhD project, and AH, AKM and RH co-supervised. TJO, AH, AKM and RH reviewed and edited the manuscript. AA updated the manuscript. All authors read and approved the final manuscript.

## Availability of data, materials and methods

All data, materials and methods are presented in this article.

